# Humanized patient-derived xenografts preserve tumour-specific immune microenvironments

**DOI:** 10.64898/2026.05.15.724697

**Authors:** Daniel Stueckmann, Jalna Meens, Jennifer Q. Pfeil, Sujana Sivapatham, Stéphane Chevrier, Shirley Hui, Christina Karamboulas, Rita Gill, Xiaoyu Zhang, Lisa Martin, Maria Komisarenko, Philip Dubé, Susan Prendeville, Hartland W. Jackson, Antonio Finelli, Gary D. Bader, Bernd Bodenmiller, Laurie Ailles, Keith A. Lawson

**Affiliations:** Department of Molecular Genetics, University of Toronto, Toronto, ON, Canada; Princess Margaret Research Institute, University Health Network, Toronto, ON, Canada; Ontario Institute of Cancer Research, Toronto, ON, Canada; Lunenfeld-Tanenbaum Research Institute, Toronto, ON, Canada; Department of Quantitative Biomedicine, University of Zurich, Zurich, Switzerland; Institute of Molecular Health Sciences, ETH Zurich, Zurich, Switzerland; Division of Urology, Department of Surgical Oncology, Princess Margaret Cancer Centre, University Health Network, Toronto, ON, Canada; Taconic Biosciences, Inc., Rensselaer, NY, USA; Department of Laboratory Medicine and Pathobiology, University Health Network, Toronto, ON, Canada; The Donnelly Centre, University of Toronto, Toronto, ON, Canada; Department of Computer Science, University of Toronto, Toronto, ON, Canada; CIFAR Multiscale Human Program, CIFAR, Toronto, ON, Canada; Department of Medical Biophysics, University of Toronto, Toronto, ON, Canada

## Abstract

Defining the genetic and cellular programs that allow solid tumours to evade immune control requires preclinical models that preserve the complexity of the human tumour immune microenvironment. Most available systems capture only part of this biology. Organoid cultures and *ex vivo* tumour fragments can retain patient-derived tumour architecture and associated immune cells, but immune populations are typically maintained only for short periods. These models also cannot capture antitumour immune responses in the physiological setting of a living organism. Patient-derived xenografts propagated in humanized mice offer a potential path to overcome these limitations by combining patient-derived tumour tissue with a reconstituted human immune system. However, few studies have systematically tested whether these models reproduce the diverse immune cell phenotypes present in the parental tumours from which they are derived. This has limited their use for studying tumour-intrinsic mechanisms that shape immune composition and promote immune evasion. To address this gap, we profiled tumour-infiltrating, splenic, and bone marrow immune cells from ovarian, head and neck, and renal PDX models propagated in CD34+ hematopoietic stem cell (HSC)-derived huNOG-EXL mice expressing human IL-3 and GM-CSF. By comparing tumours grown across distinct HSC donor backgrounds with their matched primary tumour samples, we found that tumour-intrinsic factors are a dominant determinant of immune composition in humanized PDX tumours. Across models, these immune infiltrates generally resembled those of the corresponding parental tumours. These findings support humanized PDX models as a platform for functionally interrogating tumour-intrinsic drivers of immune composition and immune evasion in solid tumours.

## Introduction

Single-cell genomic and proteomic technologies have revealed extensive heterogeneity in the immune composition of the tumour microenvironment, including marked variation among patients with the same histological diagnosis^1–4^. These studies have identified immune cell phenotypes and interactions that may be therapeutically actionable^5^. However, the field still lacks preclinical models that preserve this patient-specific biology while allowing functional testing of candidate mechanisms and targets.

*Ex vivo* tumour fragments and organoid models have helped bridge this gap by retaining patient-derived tumour architecture and, in some settings, endogenous immune populations across diverse cancer types^6–8^. However, these systems are constrained by short experimental windows, often less than 48 hours, beyond which immune-cell viability and function cannot be maintained^6^. They also lack the physiological context of an intact organism, limiting their utility for evaluating therapeutic strategies that depend on systemic immune interactions, tissue trafficking, or *in vivo* immune-cell generation and remodeling. Syngeneic and genetically engineered immunocompetent mouse models overcome some of these limitations, but they do not capture the genetic complexity of human tumours or the diversity of immune compositions observed in patients^9,10^.

Humanized immune system (HIS) mouse models provide a complementary platform for studying human tumour-immune interactions *in vivo*. These models are generated by transferring human CD34^+^ hematopoietic stem and progenitor cells (HSCs) into highly immunodeficient mouse strains, such as NSG (NOD.Cg-*Prkdc*^*scid*^ *Il2rg*^*tm1Wjl*^/SzJ) or NOG (NOD.Cg-*Prkdc*^*scid*^*Il2rg*^*tm1Sug*^Tg), where they support the development of human lymphoid and myeloid lineages^11,12^. Additional engineering to express human cytokines, including GM-CSF, IL-3, and IL-15, can improve the engraftment or differentiation of specific immune populations, such as myeloid cells or natural killer cells^13,14^. HIS mice can also support the growth of cancer cell lines and patient-derived xenografts (PDXs), offering an opportunity to study how clinically relevant genomic, transcriptomic, and metabolic features of patient tumours influence the tumour immune microenvironment^15,16^.

Despite this potential, few studies have rigorously benchmarked whether HIS models capture clinically relevant immune-cell compositions. Recent work has importantly shown that human cancer cell lines engrafted in HIS mice can generate distinct tumour immune microenvironments^16^. However, this study was limited to a small number of models, relied primarily on established cell lines rather than primary tumour fragments, used lower-resolution immune profiling, and lacked direct comparison with clinical tumour samples. It therefore remains unclear whether patient-derived tumour material engrafted into HIS mice can reproduce the immune composition of the corresponding human tumour.

Here, we used a high-dimensional cytometry by time-of-flight (CyTOF) platform to profile multilineage immune cell populations in HIS-PDX tumours, spleen, and bone marrow, and benchmarked these data against matched clinical tumour samples. Our cohort included 59 HIS-PDX mice generated using eight CD34+ HSC donors and 15 primary tumours spanning multiple histologies. This design allowed us to assess the relative contributions of HSC donor identity and tumour-intrinsic factors to both systemic immune composition and the tumour immune microenvironment. We find that immune composition within HIS-PDX tumours is primarily shaped by the engrafted tumour and, in most models, resembles the immune composition of the parental clinical tumour sample. These findings support the use of HIS-PDX models to study tumour-intrinsic determinants of human immune infiltration and immune evasion in solid tumours.

## Results

### HuNOG-EXL mice support allogenic patient derived xenograft engraftment

Solid tumours display a predominance of lymphocytes and tumour associated macrophage populations within the immune compartment^17^. To enable robust modeling of these cell types we employed the NOG-EXL strain that carries transgenes for human IL-3 and GM-CSF to enable robust myeloid cell differentiation which has historically been challenging to accomplish in HIS mouse strains^13^. We obtained eighty-eight NOG-EXL mice that were humanized using cord blood-derived CD34+ hematopoietic stem cells (HSCs) from eight donors (Taconic Biosciences), with 1 to 30 mice generated per donor (Figure 1A). Eighty-two mice were engrafted with non-HLA-matched patient-derived xenograft fragments derived from primary ovarian tumours, human papilloma virus (HPV) negative head and neck tumours, or renal cell carcinomas that had been previously established as xenografts in NSG mice (Methods). Tumour fragments were engrafted at a mean of 23 weeks (range 22-42 weeks) after humanization (Supplementary Figure 1). Six humanized mice were not engrafted with tumours to serve as controls (Figure 1B).

**Figure 1:**
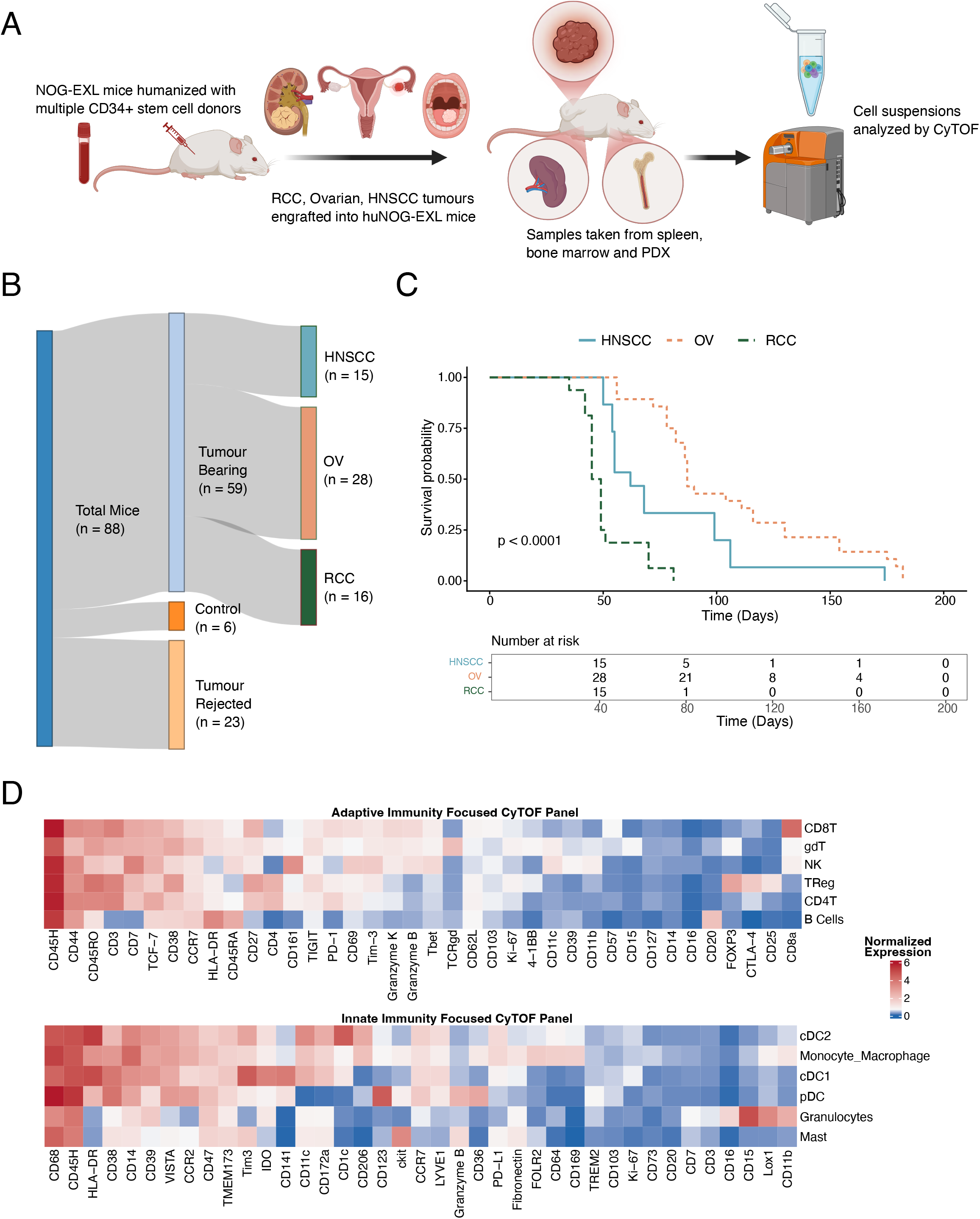
Overview of the huNOG-EXL PDX cohort. A) Schematic showing the generation of a humanized PDX cohort representing multiple stem cell donors and tumour types. B) Sankey diagram displaying the success rate for tumour engraftment in humanized mice. C) Kaplan-Meier curves displaying the survival of mice with successful tumour engraftment shown in in days following tumour implantation. Mice are grouped by tumour type (HNSCC: Head and Neck Squamous Cell Carcinoma; OV: Ovarian Cancer; RCC: Renal Cell Carcinoma. D) Heatmaps showing the average normalized expression of each marker in the adaptive immunity panel across all lymphoid cell types (top) and in the innate immunity panel across all myeloid cell types (bottom).

Tumour implantation produced 59 successful PDX tumours, corresponding to an overall engraftment rate of 72%. Engrafted models included ovarian (OV) tumours (n = 28 mice; 6 unique tumours), renal (RCC) tumours (n = 16 mice; 3 unique tumours), and head and neck (HNSCC) tumours (n = 15 mice; 4 unique tumours) (Figure 1B). Three implanted primary samples failed to engraft in any huNOG-EXL mice, and two additional primary samples lacked CyTOF data (Table 1), leaving 11 matched primary tumour-PDX pairs for downstream comparison. Although tumour implantation occurred at approximately 23 weeks after humanization, survival analysis showed that mice with successful tumour engraftment often carried tumours for several weeks to months, supporting the feasibility of extended studies in this model (Figure 1C).

**Table 1:** Description of primary tumour samples. Sixteen primary tumour samples previously established in NSG mice were implanted into huNOG-EXL mice. Passage number indicates the number of passages in NSG mice prior to cryopreservation of tumour fragments and subsequent implantation into huNOG-EXL mice.

| Manuscript Sample ID | Internal Sample ID | Sample Type | Sample Description | Passage Number | Number of PDX Replicates | Other Notes |
| --- | --- | --- | --- | --- | --- | --- |
| RCC-1 | X1-REMEDY 012-R1 | RCC tumour | Grade 3 ccRCC, pT3a | 1 | 0 | No PDX produced |
| RCC-2 | RCC-243 | RCC tumour | Grade 2 ccRCC, Stage T4NxM1 | 1 | 5 | No CyTOF data on primary (very low # of live immune cells) |
| RCC-2-CL | RCC-243-P13 | Cell line derived from RCC-2 tumour | Grade 2 ccRCC, Stage T4NxM1 | 1 | 3 | No data on primary (cell line, no immune cells) |
| RCC-3 | REMEDY005 | RCC tumour | Grade 4 ccRCC, pT3a | 1 | 8 | IMC data available for primary |
| OV-1 | X1-4424A | OV tumour | Serous ovarian cancer, collected from ascites | 1 | 3 |  |
| OV-2 | X1-6447 | OV tumour | Serous ovarian cancer, collected from ascites | 1 | 6 |  |
| OV-3 | X1-66345 | OV tumour | Serous ovarian cancer, collected from ovary | 1 | 5 |  |
| OV-4 | X1-810-1114 | OV tumour | Serous ovarian cancer, collected from pleural effusion | 1 | 4 |  |
| OV-5 | X2-3670 | OV tumour | Serous ovarian cancer, collected from omentum | 2 | 5 |  |
| OV-6 | X2-67326 | OV tumour | Serous ovarian cancer, collected from left ovary | 2 | 5 |  |
| HNSCC-1 | X1-73088 | HNSCC tumour | Stage IVA SCC, T4aN0M0. Moderate grade, collected from lower alveolus and gingiva | 1 | 0 | No PDX produced |
| HNSCC-2 | X1-73262 | HNSCC tumour | Stage II SCC, T1N1M0. Poor grade, collected from floor of mouth | 1 | 0 | No PDX produced |
| HNSCC-3 | X1-85555 | HNSCC tumour | Stage IVA SCC, T3N2bM0. Poor grade, collected from tongue | 1 | 2 |  |
| HNSCC-4 | X1-42746 | HNSCC tumour | Stage I SCC, T1N0M0. Moderate grade, collected from tongue | 1 | 4 |  |
| HNSCC-5 | X1-76489 | HNSCC tumour | Stage IVA SCC, T3N2bM0. Poor grade, collected from tongue | 1 | 4 |  |
| HNSCC-6 | X1-76695 | HNSCC tumour | Stage IVA SCC, T3N2cM0. Moderate grade, collected from tongue | 1 | 5 |  |

### PDX bearing HuNOG-EXL mice develop multilineage human immune populations

We next profiled immune cell composition across systemic and intra tumoural compartments of PDX-bearing HuNOG-EXL mice and their matched primary tumours from collected spleen, bone marrow, and tumour samples. Single-cell suspensions were analyzed by CyTOF using two previously established antibody panels^18^ enriched for adaptive and innate immune markers, respectively (Figure 1D-E), with live cells gated on human CD45 and murine CD45-H2Kd to retain human and exclude murine immune cells. Immune cells were annotated using established lineage markers (Supplementary Figure 2). A subset of lineage markers overlapping both panels enabled the annotation of six major cell populations which displayed strong concordance (Pearson correlation 0.63 - 0.96) across the innate and adaptive panels (Supplementary Figure 3).

Higher-resolution clustering revealed major lymphoid and myeloid populations (Supplementary Figures 4A-B). NK cells were defined as CD3-CD7+ and distinguished from CD3+ CD7+ T cells. T cells were further annotated as CD4+ T cells, regulatory T cells, CD8+ T cells, or γδ T cells based on expression of CD4, FOXP3, CD8, and TCRγδ. B cells were identified by CD20 expression. Dendritic cell subsets included plasmacytoid dendritic cells, conventional dendritic cell type 1, and conventional dendritic cell type 2, defined by CD123, CD141, and CD11c/CD1c expression, respectively. Mast cells were identified by c-KIT expression and separated from other CD15+ granulocytes. Monocytes and macrophages were defined as CD68+ cells lacking dendritic cell and granulocyte markers and frequently expressing CD14, CD64, and/or CD169 (Supplementary Figure 2).

Immune composition varied substantially across tissue compartments. Splenic samples had the lowest immune diversity, with a Shannon index of 1.11, and were dominated by B cells (Supplementary Figure 5A-B). PDX tumours showed the highest diversity, with a Shannon index of 1.70, and contained abundant CD4+ T cells, CD8+ T cells, regulatory T cells, cDC2s, monocytes, and macrophages (Supplementary Figure 5B). Bone marrow and primary tumour samples showed similar overall diversity, with Shannon indices of 1.67 and 1.63, respectively, but were enriched for different populations. Bone marrow contained abundant granulocytes, cDC1s, and pDCs, whereas primary tumours showed the highest proportions of monocytic cells and NK cells (Supplementary Figure 5A).

### Systemic immune composition is associated with both HSC donor and engrafted PDX

Previous studies have shown that human immune chimerism and immune composition in blood, bone marrow, and spleen are influenced by HSC donor identity in huNSG and huNSG -PDX models^15,19,20^. We therefore asked whether similar HSC donor-associated patterns were present in the bone marrow and spleen of HuNOG-EXL PDX mice. Pairwise similarity was calculated across all bone marrow samples and across all spleen samples, and sample pairs were grouped by whether they shared the same HSC donor. Bone marrow samples from mice generated with the same HSC donor had significantly more similar immune compositions than samples from different donors (median Pearson correlation 0.808 vs. 0.546, p < 0.0001; Supplementary Figure 6A). The same pattern was observed in spleen, where donor-matched samples showed greater similarity than donor-unmatched samples (median Pearson correlation 0.956 vs. 0.829, p < 0.0001; Supplementary Figure 6B).

We then repeated this analysis after grouping sample pairs by implanted tumour identity. Bone marrow and spleen samples from mice implanted with the same primary tumour were also more similar than samples from mice implanted with different tumours (bone marrow: median Pearson correlation 0.831 vs. 0.610, p < 0.0001; spleen: median Pearson correlation 0.935 vs. 0.852, p < 0.0001; Supplementary Figure 6C-D). These data suggest that systemic immune composition in PDX bearing huNOG-EXL mice is shaped by both HSC donor identity and the implanted tumour.

### Intratumoural immune composition is driven by engrafted PDX

Within each tumour type, we observed both highly infiltrated models, including RCC-3, OV-3, and HNSCC-4, and poorly infiltrated models, including RCC-2, OV-4, and HNSCC-5 which were consistent across PDX humanized with different HSC donors (Supplementary Figures 5C-E). HSC donor identity was associated with variation in peripheral blood chimerism measured 10 weeks after humanization (Supplementary Figure 7A), but blood chimerism did not correlate with subsequent human immune infiltration into PDX tumours (Pearson correlation = -0.217; Supplementary Figure 7B). Together, these analyses indicate that the extent of immune infiltration in PDX tumours is tumour dependent.

We next tested whether the composition of the tumour immune microenvironment was also tumour driven. To compare PDX tumours with different levels of infiltration, each immune population was normalized to the total human immune infiltrate within each sample. PDX tumours were then clustered according to immune cell composition (Figure 2A). Replicate PDX tumours derived from the same primary sample showed reproducible immune compositions and clustered by primary tumour identity rather than HSC donor identity. Pairwise Pearson correlation confirmed that PDX tumours derived from the same primary tumour were significantly more similar to one another than PDX tumours derived from different primary tumours (median Pearson correlation 0.909 vs. 0.537, p < 0.0001; Figure 2B). In contrast, PDX tumours generated with the same HSC donor were no more similar to one another than tumours generated with different donors (median Pearson correlation 0.560 vs. 0.558, p = 0.93; Figure 2C). These findings indicate that both the extent and composition of immune infiltration in huNOG-EXL PDX tumours are primarily shaped by the implanted tumour rather than by the HSC donor.

**Figure 2:**
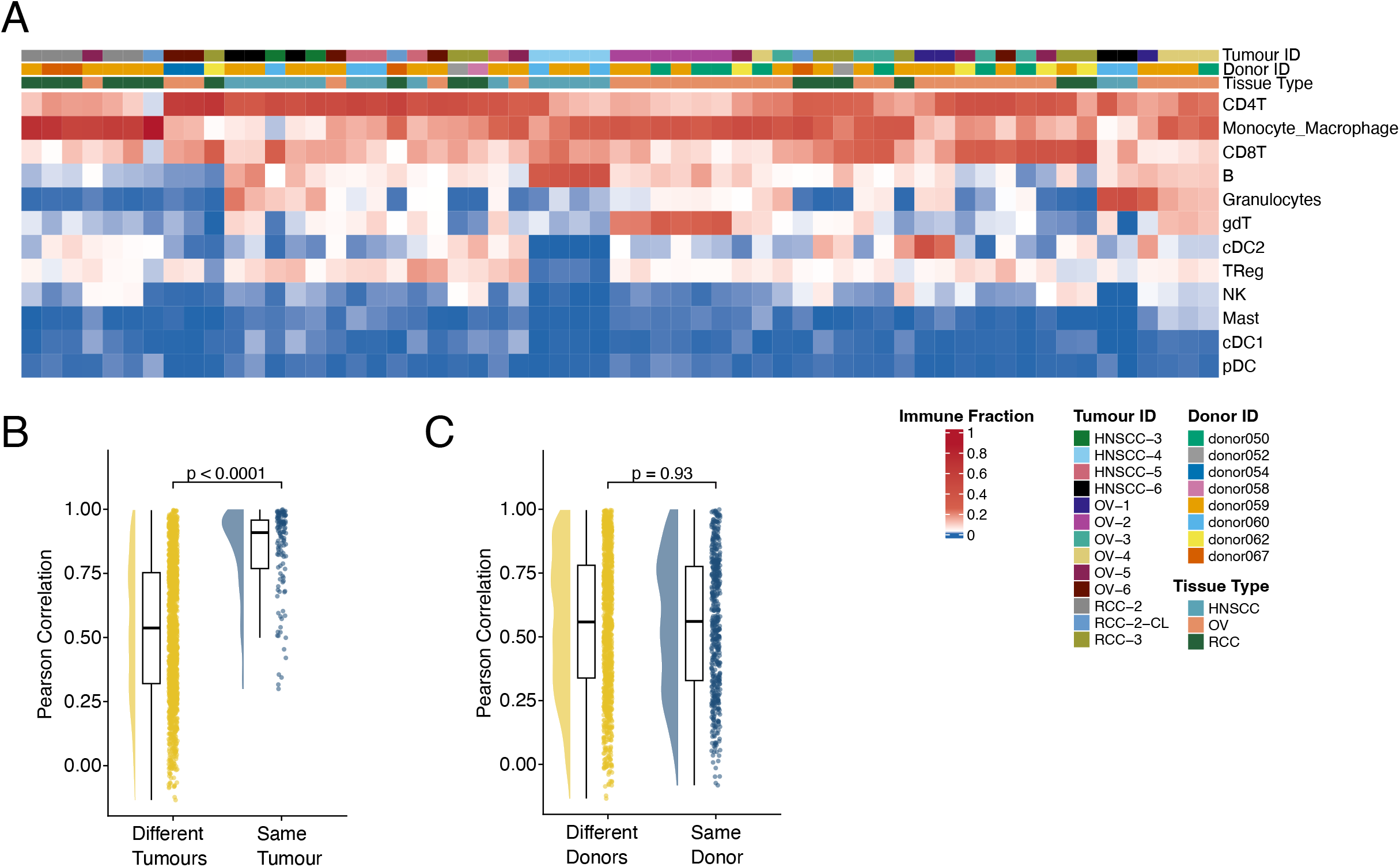
Tumour-intrinsic factors drive immune composition in humanized PDX tumours. A) Heatmap displaying the abundance of each immune cell type as a proportion of total human immune cells within each PDX sample (n = 59). The CD34+ stem cell donor ID used to humanize each mouse, the primary tumour ID implanted into each mouse, and the tumour type are shown for each PDX. B/C) Violin plots showing the distribution of pairwise comparisons of the similarity of PDX immune composition. Similarity was quantified as the pearson correlation of immune cell type proportions between all possible pairs of PDX samples shown in (A). Pairwise comparisons were grouped into two distributions using metadata shown in (A), and the distributions were compared using Student’s t-test. Pairs of PDX were classified as having the same (blue) or different (yellow) tumour IDs (B), or the same (blue) or different (yellow) stem cell donor IDs (C). Pairs of PDX implanted with the same primary tumour showed signficantly greater similarity than pairs of PDX with different implanted tumours. Dislayed p values represent results of two-sided Student’s t-tests.

### Imaging mass cytometry validates tumour-driven PDX similarity

To validate the tumour-driven organization of the PDX immune microenvironment using an orthogonal spatial platform, we analyzed 14 renal cancer PDX tumours by imaging mass cytometry (IMC) using a custom antibody panel (Table 2). This approach allowed simultaneous quantification of immune, stromal, endothelial, and cancer cells. Marker-based annotation identified CA9+ cancer cells as the dominant population, along with smaller populations of PECAM1+ endothelial cells and SMA+ stromal cells (Supplementary Figure 8A). Consistent with the CyTOF data, the most abundant immune populations in renal PDX tumours were monocytes and macrophages and CD4+ T cells (Supplementary Figure 9A).

**Table 2:** Custom antibody panel for IMC acquisition. Antibody targets are shown alongside their conjugated heavy metal isotopes.

| Metal Tag | Target | Cell Type/State | Antibody Clone | Lot |
| --- | --- | --- | --- | --- |
| In113 | E-Cad / P-Cad | Epithelial | 36/E-Cadherin | 2287671 |
| In115 | (CK2) pan Cytokeratin | Epithelial | AE1 | YF3946143 |
| In115 | Keratin Epithelial | Epithelial | AE3 | 3460341 |
| In115 | (CK1) pan Cytokeratin | Epithelial | C11 | 3382323 |
| Pr141 | HLA-DR | MHCII | TAL 1B5 | 1011954-1 |
| Nd143 | PD-L1 | Exhaustion | CAL10 | GR3428776-3 |
| Nd144 | Cleaved Caspase3 | Apoptosis | Polyclonal_CST9661 | 48 |
| Nd145 | CD15 | Granulocytes | EPR22011-11 | B265372 |
| Nd146 | CD45RA | Naïve | HI100 | 2181694 |
| Sm147 | CD163 | Macrophages | EDHu-1 | 149022A |
| Nd148 | CD16 | Non-classical monocytes | EPR16784 | GR3416046-5 |
| Sm149 | CD20 | B cells | L26 | 2526316 |
| Nd150 | CD68 | Pan myeloid | KP1 | B283618 |
| Eu151 | CD4 | Helper T cell | EPR6855 | 1023375-4 |
| Sm152 | CD8a | Cytotoxic T cell | C8/144B | B369894 |
| Sm154 | CD11c | Dendritic cells | EP1347Y | GR3414808-5 |
| Gd155 | mitochondria | human cells | 113-1 | 1012031-1 |
| Gd156 | FOXP3 | Treg | 221D |  |
| Gd157 | TREM2 | Macrophages | D8I4C | 1 |
| Gd158 | LYVE-1 | Macrophages | RM1008 | GR3399057-5 |
| Tb159 | goat IgG (H+L) | secondary |  | YF3974659D |
| Tb159<br>Secondary | CXCL10 | Macrophages | Polyclonal_AF-266-NA | ABR1722071 |
| Gd160 | FAP | Fibroblasts | Polyclonal_AF3715 | ZKWO622011 |
| Dy161 | CAIX | RCC | polyclonal_CA9_AF2188 | VNQ0319071 |
| Dy162 | CD45RO | Memory | UCHL1 | 2 |
| Dy163 | Rat IgG | secondary | Polyclonal_A18873_Thermo | 61-172-060320 |
| Dy163<br>Secondary | CD3 | pan T cells | CD3-12 | YE3922415 |
| Dy164 | Granzyme B | Effector | D6E9W | 2 |
| Ho165 | CTLA-4 | Exhaustion | CAL49 | 1045701-1 |
| Er166 | CXCL13 | TLSs | Polyclonal_CXCL13 / BLC / BCA-1 | BAS0317111 |
| Er167 | FOLR2 | Macrophages | EPR25731-70 | GR3457571-1 |
| Er168 | Ki-67 | Proliferation | B56 | 3026625 |
| Tm169 | Mouse IgG (H+L) | secondary | Polyclonal_A28174_Thermo | 2276480 |
| Tm169<br>Secondary | TCR gamma/delta | $\gamma\delta$ T cells | H-41 | H2321 |
| Er170 | CD117 (c-kit) | Mast | D3W6Y | 1 |
| Yb171 | TCF7 | secondary | C63D9 | RL246119A |
| Yb172 | CD14 | Monocytes | SP192 | 2 |
| Yb173 | CD11b | Natural killer cells |  | GR3438520-5 |
| Yb174 | CD45 | Pan immune cell | D9M8I | 2 |
| Lu175 | CD1c | cDC2 | OT12F4 | F004 |
| Yb176 | CD279 (PD-1) | Exhaustion | D4W2J | 7 |
| Pt195 | PNA<sub>d</sub> | TLSs | MECA-79 | B341880 |
| Pt196 | CD31 | Endothelial | RM1006 | GR3438767-1 |
| Y89 | SMA | Pericytes | 1A4 | 2382312 |
| Bi209 | rabbit IgG | secondary |  |  |
| Y89<br>Secondary | CD45 mouse | mouse immune cells |  |  |

We then calculated pairwise similarity across 69 IMC regions from the 14 renal PDX tumours. Regions from PDX tumours derived from the same primary tumour were more similar than regions from different primary tumours (median Pearson correlation 0.863 vs 0.802, p < 0.0001; Supplementary Figure 9B). This subset of the PDX cohort represented a limited number of combinations of HSC donor background and implanted tumour and was therefore underpowered to validate the impact of HSC donor on immune composition; however, these spatial data support the CyTOF-derived findings that tumour-intrinsic factors shape the intratumoural immune cell composition.

### PDX tumours contain heterogeneous T cell and macrophage states

Having established that immune cell abundance is tumour dependent, we next asked whether immune cell phenotypes within major lineages showed similar tumour-intrinsic patterns. We focused first on CD8+ T cells, which can occupy distinct functional states across a spectrum from cytotoxic (marked by granzyme B and granzyme K) to exhausted (marked by Programmed cell death protein 1 (PD-1) and T cell immunoglobulin and mucin domain-containing protein 3 (TIM-3)).

CD8+ T cell marker expression varied across PDX models (Figure 3A, top), and there were apparent patterns of tumour-specific marker expression spanning multiple HSC donor backgrounds. Four PDX tumours derived from HNSCC-4 across two HSC donor backgrounds showed strong granzyme B expression and moderate TIM-3 and granzyme K expression. Two PDX tumours derived from OV-3 showed high expression of both TIM-3 and PD-1, consistent with a more exhausted phenotype. Euclidean distance analysis showed that CD8+ T cell expression profiles were more similar among PDX tumours derived from the same primary sample than among tumours derived from different primary samples (p < 0.0001; Figure 3B). These data suggest that PDX tumours can reproducibly capture tumour-associated CD8+ T cell phenotypes.

**Figure 3:**
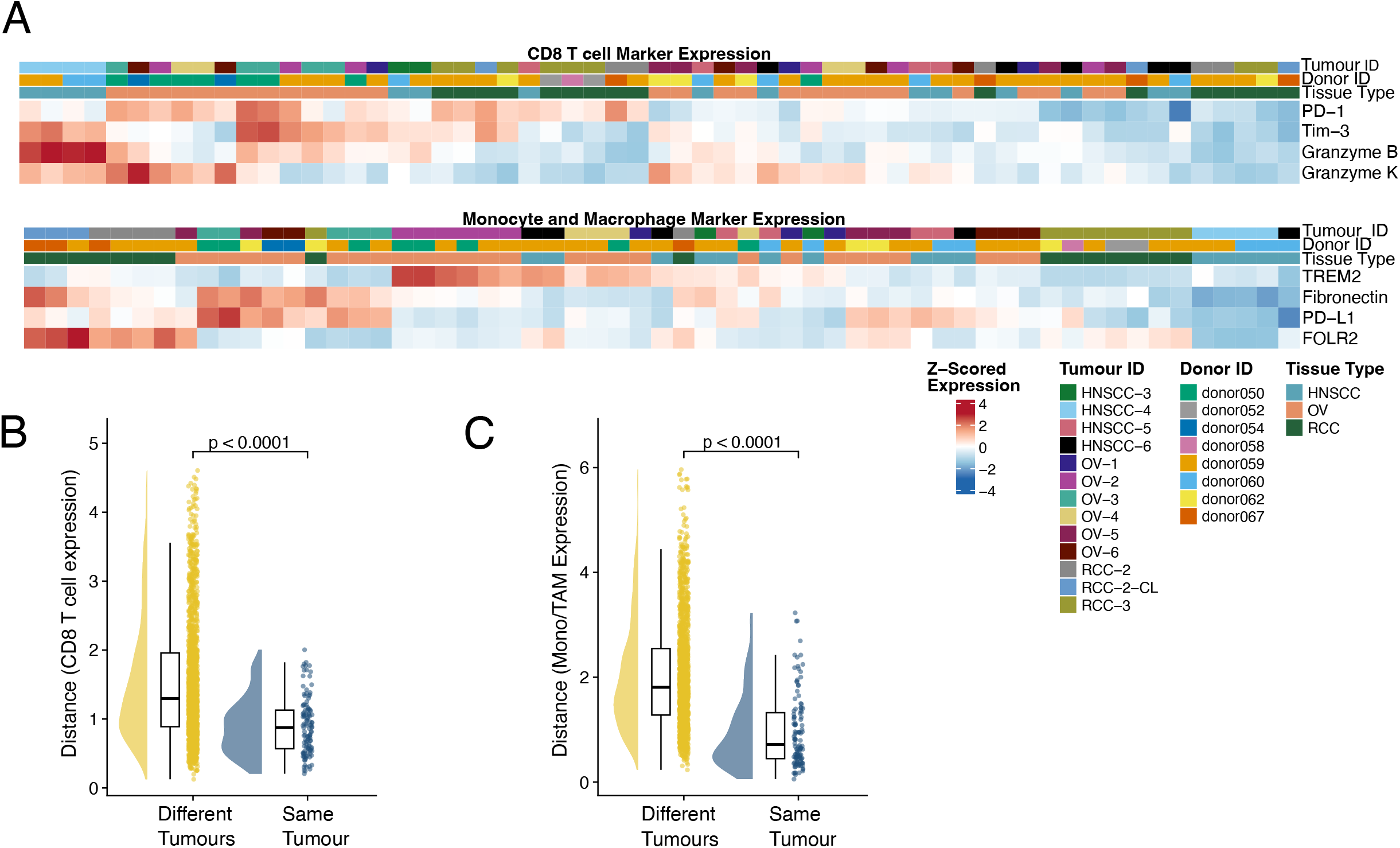
Tumour-intrinsic factors drive immune cell state in humanized PDX tumours. A) Heatmap displaying the z-scored average expression of four CD8+ T cell state markers within annotated CD8+ T cells (top) and the z-scored average expression of four macrophage subtype markers within annotated monocytes and macrophages (bottom) within each PDX B) Rain plots showing the distribution of pairwise comparisons of the similarity of CD8+ T cell expression profiles. Similarity was quantified as the euclidean distance between all possible pairs of PDX samples shown in the CD8 T cell expression heatmap (A). Pairwise comparisons were grouped into two distributions on the basis of having the same tumour ID (blue) or different tumour IDs (yellow), and the distributions were compared using Student’s t-test with two-sided p values shown. C) Rain plots showing the distribution of pairwise comparisons of the similarity of monocyte and macrophage expression profiles. Similarity was quantified as in (B) between all possible pairs of PDX samples shown in the monocytic expression heatmap (A). Pairwise comparisons were grouped into two distributions as in (B), and the distributions were compared using Student’s t-test.

We performed a parallel analysis of monocytes and macrophages, which also comprise heterogeneous states in human tumours. Monocytes and macrophages were assessed for expression of fibronectin, Triggering Receptor Expressed on Myeloid cells 2 (TREM2), Folate Receptor Beta (FOLR2), and Programmed death-ligand 1 (PD-L1). This analysis identified distinct groups of PDX tumours, including a cluster composed mainly of renal PDX tumours with high FOLR2 expression representing multiple HSC donor backgrounds (Figure 3A, bottom). As observed for CD8+ T cells, monocyte and macrophage expression profiles were more similar among PDX tumours derived from the same primary sample than among tumours derived from different primary samples (p < 0.0001 Figure 3C). Thus, tumour-associated patterns extend beyond immune-cell abundance to include lineage-specific phenotypic states.

### Cell line-derived xenografts resemble matched PDX tumours

Cancer cell lines derived from patient tumours are widely used pre-clinical models because they are easier to expand, store, and manipulate than primary tumour fragments. To determine whether patient-derived cell lines generate tumour-specific immune microenvironments in huNOG-EXL mice, we compared PDX and cell line-derived xenograft (CDX) tumours established from the same primary renal tumour. Three mice were implanted with RCC-2-CL, a patient-derived cell line generated from the RCC-2 tumour. These CDX tumours contained immune infiltrates dominated by monocytes and macrophages, with substantial CD4+ T cell infiltration, a pattern similar to that observed in five matched RCC-2 PDX tumours (Figure 4A).

**Figure 4:**
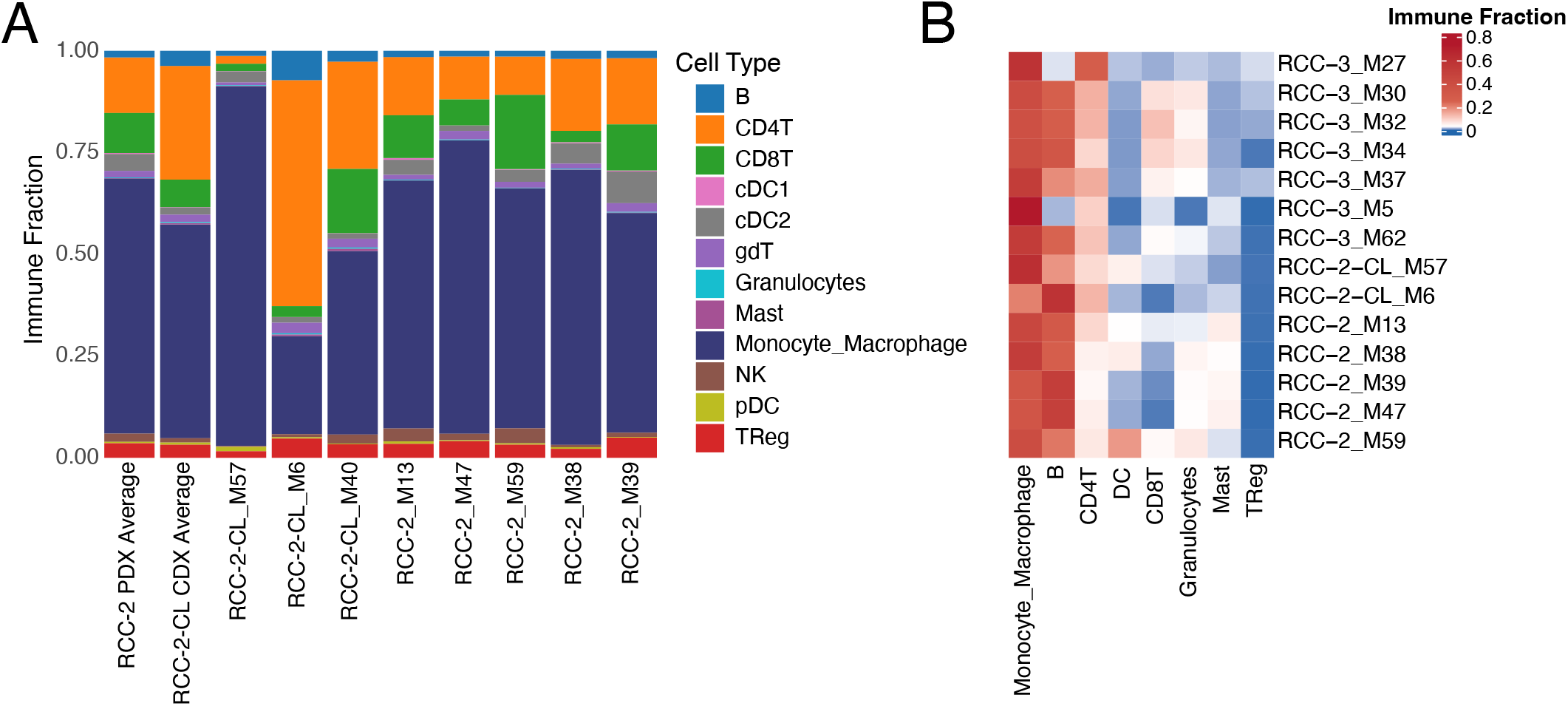
Cell line-derived xenografts display similar immune composition to matched PDXs. A) Stacked bar plots displaying the proportion of all immune cell types within PDX and CDX generated from a primary RCC tumour (RCC-2). Five PDX were produced using this primary tumour (RCC-2), and three CDX were generated from a cell line derived from RCC-2 (RCC-2-CL). Average immune composition across the five PDX and three CDX are shown as “PDX Average” and “CDX Average” respectively. All PDX and CDX derived from RCC-2 displayed extensive myeloid and CD4+ T cell infiltration with low CD8+ T cell infiltration. B) Heatmap displaying the proportion of immune cell types captured using IMC data in RCC PDX and CDX including 7 of the 8 samples shown in (A).

IMC analysis further supported the similarity between RCC-2 PDX and RCC-2-CL CDX tumours. Both groups showed reproducible immune compositions that differed from RCC-3 PDX tumours (Figure 4B). All RCC tumours displayed high monocyte/macrophage and CD4+ T cell infiltration. However, RCC-2 and RCC-2-CL tumours exhibited infiltration of dendritic cells and mast cells, whereas RCC-3 tumours contained higher proportions of CD8+ T cells and regulatory T cells. These findings confirm that in this renal cancer example, xenografts generated from a patient -derived cell line showed immune compositions similar to those observed in xenografts established from the original tumour.

### PDX tumours recapitulate key features of matched primary tumour immune composition

As PDX tumours showed variable levels of human immune infiltration, we examined whether this variation resembled the heterogeneity of the primary tumour samples. Immune infiltration differed by tumour type, with ovarian primary tumours and ovarian PDX tumours showing low overall immune infiltration (Supplementary Figure 5F). Across the cohort, average immune infiltration was lower in PDX tumours than in primary tumour samples, although this difference did not reach statistical significance (mean human immune infiltration 15.9% vs 37.8%, p = 0.12; Supplementary Figure 5F). This trend may in part reflect the composition of the cohort, as nearly half of the PDX tumours were derived from ovarian cancers.

The reproducible, tumour-driven composition of PDX immune microenvironments prompted us to compare PDX tumours directly with their matched parental primary tumour samples. Primary tumour cell suspensions were profiled using the same CyTOF workflow as the mouse -derived samples. PDX tumours were classified as matched if they originated from the same primary tumour and unmatched if they originated from a different primary tumour. Two primary samples had insufficient immune cell numbers and were excluded, leaving 11 primary tumours with at least one matched PDX tumour.

Seven of the 11 primary tumours were more similar to their matched PDX tumours than to unmatched PDX tumours (Figure 5A). Across the cohort, primary tumours were significantly more similar to matched PDX tumours than to unmatched PDX tumours (median Pearson correlation of 0.628 vs. 0.520, p = 0.012; Figure 5B). Models that did not closely resemble their matched primary tumours tended to show the largest reductions in monocytes and macrophages, suggesting that myeloid representation remains incomplete even in the myeloid-supportive huNOG-EXL strain (Supplementary Figure 10A-B). This was most evident in PDX tumours derived from HNSCC-5 and OV-6. The HNSCC-5 primary tumour was heavily infiltrated by monocytic cells, which comprised approximately 60% of the immune compartment, whereas matched HNSCC-5 PDX tumours contained approximately 10% monocytic cells (Supplementary Figure 10A). A similar pattern was observed for OV -6, in which monocytic cells comprised approximately 80% of the primary tumour immune compartment but only approximately 10% of the immune infiltrate in each of four matched PDX tumours (Supplementary Figure 10B).

**Figure 5:**
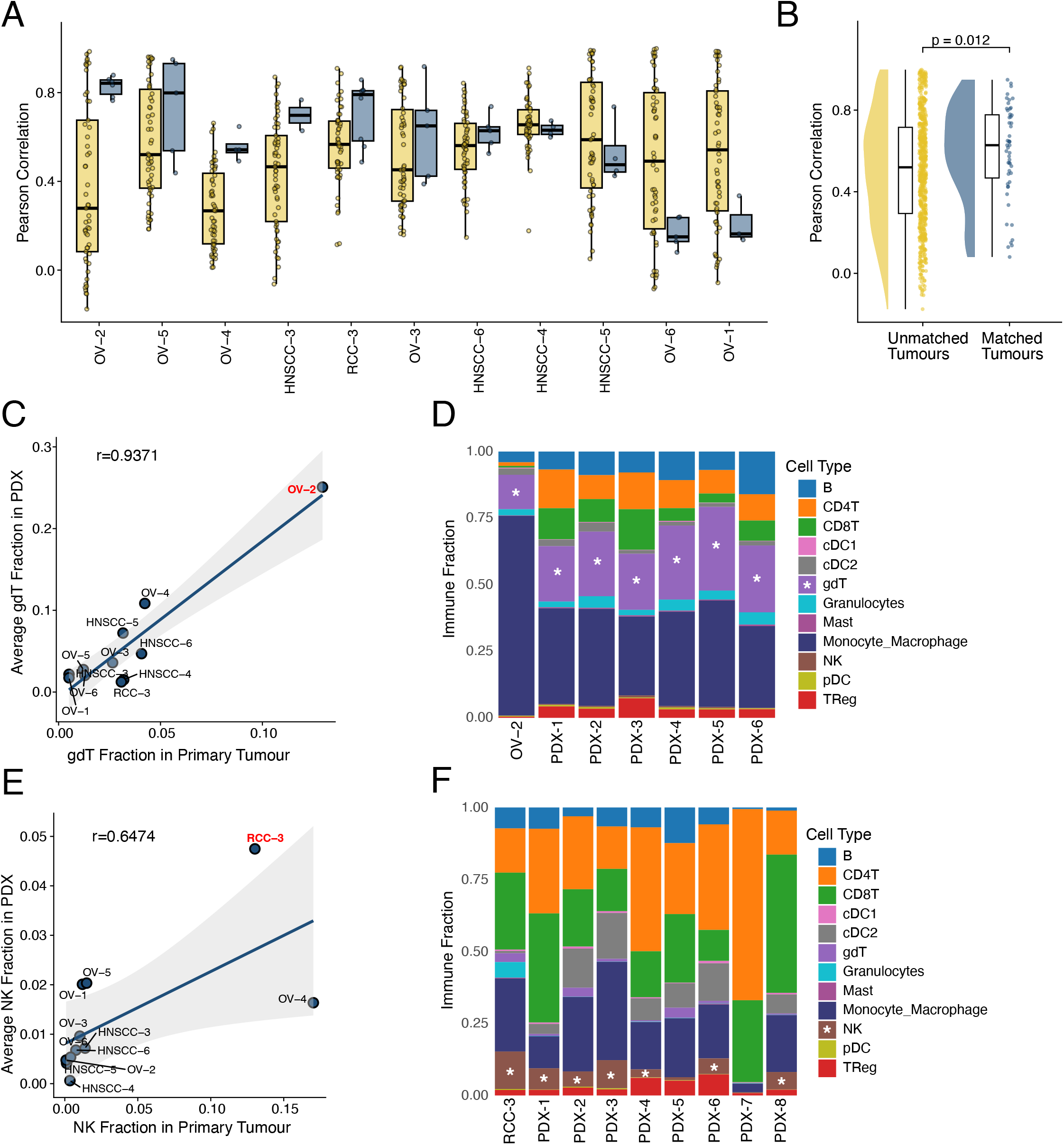
PDX tumours recapitulate key features of matched primary tumour immune composition. A) Barplots displaying the pairwise similarity between the immune composition of n = 11 primary tumour samples to all n = 59 PDX. Similarity was quantified as the Pearson correlation of immune cell type proportions between all possible combinations of primary tumour samples and PDX, and is displayed for each primary tumour. Primary tumour samples which were not successfully engrafted into any mice were excluded from this analysis. Pairwise comparisons were grouped into two distributions on the basis of having the same tumour ID (blue) or different tumour IDs (yellow). Seven of the 11 primary tumours displayed greater similarity to tumour-matched PDX than unmatched PDX. B) All data from (A) were summarized into two distributions and were compared using Student’s t-test. C) A scatter plot displaying the infiltration of gamma-delta T cells in all n = 11 primary tumour samples and PDX derived from these samples. The proportion of gamma-delta T cells was averaged across all PDX replicates originating from the same primary tumour. An ovarian sample (OV-2) is highlighted as having high gamma-delta T cell infiltration in both the primary and PDX samples. D) Stacked bar plots showing the proportion of all cell types in the OV-2 primary sample and all six PDX produced from OV-2 (labelled as PDX # 1 to 6). E) A scatter plot displaying the infiltration of NK cells in all n = 11 primary tumour samples and PDX derived from these samples. The proportion of NK cells was averaged across all PDX replicates originating from the same primary tumour. An RCC sample (RCC-3) is highlighted as having high NK cell infiltration in both the primary and PDX samples. F) Stacked bar plots showing the proportion of all cell types in the RCC-3 primary sample and all eight PDX produced from RCC-3 (labelled as PDX # 1 to 8). Stars indicate populations of interest. Correlation (r) values shown are Pearson correlation coefficients.

We also identified primary tumours with distinctive immune features that were preserved in matched PDX tumours. Despite being a rarer population (Supplementary Figure 5A), the intratumoural fraction of γδ T cells showed strong concordance between primary tumours and matched PDX samples (Pearson correlation = 0.9371; Figure 5C). The OV-2 primary tumour notably contained abundant γδ T cells, and all six matched OV-2 PDX tumours generated across two HSC donors contained γδ T cells representing more than 20% of the immune infiltrate (Figure 5D).

Similarly, NK cells appeared at low frequencies in most tumour samples (Supplementary Figure 5A) but displayed concordance between primary tumours and matched PDX samples (Pearson correlation = 0.6474; Figure 5E). The RCC-3 primary tumour showed extensive CD4+ T cell, CD8+ T cell, and NK cell infiltration (Figure 5F). Eight matched RCC-3 PDX tumours generated across four HSC donors exhibited strong CD4+ and CD8+ T cell infiltration, and six of eight also displayed NK cell infiltrates (Figure 5F). These examples show that huNOG-EXL PDX tumours can preserve distinctive immune features of their parental tumours.

Because tissue dissociation can affect recovery of specific immune populations in single-cell assays^21^, we used IMC to validate the presence of NK cell infiltration in RCC-3 PDX tumours. Tissue sections from multiple RCC-3-derived PDX tumours were analyzed by IMC (Supplementary Figure 8B). In addition to CA9+ tumour cells and SMA+ stromal cells, RCC-3 PDX sections contained numerous CD45+ CD3+ T cells. We also identified NK cell populations marked by CD11b expression with lack of CD68 and CD14 expression (Supplementary Figure 8B). These spatial data support the CyTOF results and confirm that RCC-3 PDX tumours contain both T cell and NK cell infiltrates.

## Discussion

Humanized immune system PDX models provide an opportunity to study patient-derived tumour-immune interactions *in vivo*, but their fidelity to clinical tumour immune microenvironments has not been systematically evaluated. Here, we show that huNOG-EXL PDX models generate reproducible tumour immune microenvironments across HSC donors. In some cases, matched PDX tumours retained distinctive immune features of the parental clinical tumour, including γδ T cell enrichment in an ovarian cancer model and NK cell infiltration in a renal cancer model. These data support huNOG-EXL PDX models as a platform for studying patient-specific features of tumour immune composition and for functionally testing tumour-intrinsic mechanisms that shape immune infiltration.

The durability of the HuNOG-EXL model further supports its use for preclinical immunotherapy studies. In contrast to reports of more limited experimental windows in NSG-SGM3 mice^22^, huNOG-EXL mice supported tumour implantation more than 20 weeks after humanization, allowing tumours to be established after multilineage immune differentiation. In this study, implanted tumours were routinely followed for more than 50 days before reaching endpoint criteria. This extended window should accommodate many immunotherapy study designs, including approaches that require repeated dosing, longitudinal sampling, or delayed immune remodeling.

Consistent with prior work, huNOG-EXL mice showed HSC donor-dependent differences in systemic immune composition and early blood chimerism^15,20^. However, these donor effects did not explain the level or composition of immune infiltration within PDX tumours. Instead, tumour identity was the dominant determinant of immune infiltration and immune composition within the tumour microenvironment, often capturing key features of the matched clinical human tumour. This finding provides a foundation for using these models to discover clinically relevant tumour-intrinsic genetic and cellular drivers of immune evasion, representing a key direction of future work. We also anticipate that these models will enable direct functional testing of strategies that target specific immunosuppressive populations, including heterogeneous tumour-associated myeloid populations for which antibody-based, CAR T-cell-mediated, and reprogramming approaches are being developed^23–25^.

### Limitations of the study

Solid tumours vary widely in mutational burden, driver alterations, transcriptional and metabolic states, all of which have been linked to immune infiltration^26–30^. These features are likely to contribute individually or synergistically to drive the differences in immune microenvironments observed across tumour types and patient samples in this study. Hence, identifying the specific tumour-intrinsic factors that drive our observed immune cell infiltration and composition within HIS-PDX models will require additional molecular profiling and functional validation.

CyTOF allowed us to quantify more than 60 proteins at single-cell resolution and identify major immune populations across tumour, spleen, and bone marrow samples. However, this approach has lower molecular depth than single-cell RNA sequencing or spatial transcriptomic methods^31^. As a result, closely related cell states such as B cells and plasma cells could not be resolved with confidence despite expected differences in CD20 and CD38 expression^18^, possibly due to low plasma cell abundance in this model^32^. Similarly, monocytes and macrophages formed a continuum of marker expression rather than discrete populations, limiting our ability to resolve finer myeloid states. Lastly, preservation of spatially organized immune cell networks, such as tertiary lymphoid structures, were not systematically assessed.

Single-cell and spatial atlases of human tumours have revealed extensive heterogeneity in myeloid and T cell states, as well as in their spatial organization, across patients and cancer types^1–3,33^. Although our data show that HIS-PDX models capture patient-specific immune compositions, some aspects of the immune microenvironment remain incompletely represented, including tumour-associated myeloid populations in certain tumour models. Beyond these limitations, allogeneic humanized mouse models remain constrained in their ability to model antigen-specific adaptive immunity^34^. Further work will be required to benchmark models engineered to support lymphocyte education by human thymic tissue^34,35^.

## Acknowledgements

A.F. and H.W. J. were supported by the Office of the Assistant Secretary of Defense for Health Affairs through the Kidney Cancer Research Program Translational Research Partnership Award (Award No. W81XWH2210868). The opinions, interpretations, conclusions, and recommendations expressed herein are those of the authors and are not necessarily endorsed by the Department of Defense.

K.A.L and A.F. were also supported by the Princess Margaret Cancer Foundation, the Canadian Urological Association Scholarship Foundation (CUASF), and the Canadian Urological Association in collaboration with the Kidney Cancer Research Network of Canada (KCRNC).

L.A. is supported by an Ontario Institute for Cancer Research Investigator Award (OICR IA -016). K.A.L is supported by a Hold Em for Life Early Career Professorship in Cancer Research.

Schematic figures were created with BioRender.com.

## Declaration of interests

Taconic Biosciences Inc provided the mice used in this study under a sponsored research agreement. K.A.L. receives reports receiving honorarium from AbbVie and speaking fees from Virtual Hallway. H.W.J has consulted for and received travel and research support from Standard BioTools unrelated to this work. G.D.B. has equity in Adela Inc. and is an advisor of BioRender. B.B. has founded and is a shareholder and member of the board of Navignostics, a precision oncology spin-off from the University of Zurich. P.D. is an employee of Taconic Biosciences.

## Author contributions

Conceptualization, L.A, K.A.L; Methodology, J.M, J.Q.P, S.S, C.K, R.G, X.Z; Formal analysis, D.S, S.S, S.C, S.H, S.P; Writing - D.S, K.A.L; Writing - Review & Editing, All authors; Supervision, P.D, H.W. J, G.D.B, A.F, B.B, L.A, K.A.L; Funding Acquisition, H. W. J, A.F, L.A, K.A.L. All authors reviewed and approved the manuscript.

## Declaration of AI-assisted technologies in manuscript preparation

The authors used ChatGPT (OpenAI) and Claude to edit for grammar, clarity, and flow. The authors reviewed, revised, and approved the final text and take full responsibility for the content, interpretation, and conclusions presented.

## Data and code availability

### Data

Raw data and preprocessed data objects will be made available upon request.

### Code

All code will be made publicly available at https://github.com/lawsonlab-uhn/humanized-PDX; the repository is currently private and will be made available upon publication.

## Experimental Procedures

### Patients and Tissue Samples

Tumour samples were obtained from patients undergoing surgery at the University Health Network. All samples were collected from patients with informed consent, and all related procedures were performed with the approval of the Research Ethics Board (REB# 12-5639, REB# 19-6304, REB# 09-0828-T, REB# 17-5204, REB# 21-5487, REB# 18-5842, REB# 22-5377) of the University Health Network. Patient samples were dissociated into single cells and viably frozen into single cell suspensions for further testing at a later time, as previously described^33^. Briefly, tumours were manually cut into small pieces and incubated at 37 degrees in a cocktail of M199 + 1X Collagenase/Hyaluronidase (Stem Cell Technologies) + DNase (Worthington) and incubated for a period of 2 -3 hours with manual trituration. Samples were then filtered through a 60 micron mesh and red blood cells were lysed in ACK lysis buffer (Thermo Fisher). Samples were then viably frozen in 90% FBS (GIbco) + 10% dimethylsulfoxide (DMSO).

### Initial generation of PDX models

All animal experiments were performed with the approval of the University Health Network Animal Care Committee and adhered to the Canadian Council on Animal Care guidelines (protocol #1542). NOD/SCID/IL2Rγ^−/−^ (NSG) mice were bred in-house at the University Health Network Animal Resources Centre. Patient Tumour tissues were dissected into small fragments (∼ 1mm^3^) and five to ten fragments per patient were implanted into of NSG mice (maximum two fragments per mouse). HNSCC tumours were implanted under the skin on the flanks of NSG mice, RCC tumours were implanted into the renal capsule, and OV tumours were implanted in the mammary fat pad. Mice were monitored weekly for tumour growth and were euthanized when tumours reached an end point of 1.5 cm diameter, or at 6 months post-implantation. Tumour fragments from established PDXs were viably banked by cryopreservation in 90% fetal bovine serum (FBS)/10% Dimethyl Sulfoxide (DMSO) or dissociated into single cell suspensions and viably frozen as described previously^36^.

### Generation of huNOG-EXL PDX

Female NOD.Cg-*Prkdc*^*scid*^*Il2rg*^*tm1Sug*^Tg(SV40/HTLV -Il3,CSF2)10-7Jic/JicTac mice engrafted with human cord blood HSCs (huNOG-EXL) were provided by Taconic at 16-17 weeks post-engraftment. PDX models were divided into groups of 6 mice split between 2 donors with similar expression of CD45 percentage populations. Implantation of HNSCC and RCC samples were done by either surgical implantation of previously generated cryopreserved PDX tissue fragments, placed subcutaneously in the left flank or by injecting dissociated PDX tumours subcutaneously 1:1 with Matrigel in the left flank as previously described^37^. OV PDX cell suspension samples were injected into the number five mammary fat pad with a 1:1 ratio of cells to Matrigel. Mice were monitored weekly for tumour growth and overall health status and tumours were harvested at a maximum diameter of 1.5cm, or previously outlined humane endpoint in our Animal User Protocol (protocol #1542). Several mice were noted to suffer from hair loss and skin rashes which were noted at the time of euthanasia and suspected to be caused by Graft vs Host Disease. All animal experiments were performed with the approval of the University Health Network Animal Care Committee and adhered to the Canadian Council on Animal Care guidelines.

### Sample Processing

Tumour samples and spleens were collected at endpoint and dissociated into single cell suspensions and viably frozen, as previously described^38^. Bone marrow was collected by removal of the intact femur bones removing all muscle and tissue from the bones on both sides of the mouse. A 26g needle and syringe containing 5 milliliters of M199 media was inserted into the bone cavity via the patellar surface. The head of the femur was cut off and the bone marrow was flushed into a collection tube. The samples were spun, filtered using a 60um mesh followed by a red blood cell lysis step then washed once in complete IMDM media containing 10% FBS, and viably frozen in 90% FBS containing 10% DMSO.

### Antibodies and Antibody Labelling

Antibodies used in suspension and imaging mass cytometry analyses were conjugated using the MaxPAR antibody labeling kit (Fluidigm). Following conjugation, antibody recovery yield was quantified by Nanodrop spectrophotometry (Thermo Scientific), and the conjugated antibodies were stored at 4°C with Candor Antibody Stabilizer. Optimal working concentrations for all conjugated antibodies were determined empirically through titration experiments. Antibody inventory and metadata were maintained using the cloud-based laboratory management platform AirLab^35^. Two panels of 40 antibodies each were used to analyze cell suspensions from primary tumour, PDX, bone marrow and spleen samples. In addition to cell lineage and state markers (Figure 1D-E), the panels contained antibodies for cleaved caspase 3/PARP to detect apoptotic cells and cisplatin for dead cells. Murine immune cells were identified with CD45M-H2Kd double positive staining to differentiate from human immune cells.

### CyTOF Data Acquisition

Viably frozen cell aliquots were thawed and resuscitated according to standard protocols. Following centrifugation, cells were washed with RPMI1640 medium and re-centrifuged. Aliquots of up to 1 × 10^7 cells were resuspended in 1uM cisplatin Pt194 (Fluidigm) working stock diluted in RPMI1640 and incubated for 15min at 4°C. The staining was quenched with 1 mL cell staining medium (CSM: PBS with 0.5% bovine serum albumin and 0.02mM EDTA). Cells were centrifuged and resuspended in 1.6% paraformaldehyde (PFA, Electron Microscopy Sciences) diluted in RPMI1640, and fixed at room temperature for 10 min. The fixation reaction was quenched by adding 1 mL of CSM. Cells were centrifuged, and the disrupted pellet was stored at −80°C until the debarcoding step.

To ensure homogeneous staining across samples, we employed a mass tag barcoding strategy using 1 × 10^6 cells per sample. A 60 -well barcoding scheme consisting of unique combinations of four out of eight barcoding reagents was used as previously described^39^. Six palladium isotopes (102Pd, 104Pd, 105Pd, 106Pd, 108Pd, and 110Pd, Fluidigm) were chelated to 1-(4-isothiocyanatobenzyl)ethylenediamine-N,N,N’,N’-tetraacetic acid (Dojino), and two indium isotopes (113In and 115In, Fluidigm) were chelated to 1,4,7,10-tetraazacyclododecane-1,4,7-tris-acetic acid 10-maleimide ethylacetamide (Dojino) following standard procedures^40^. Barcoding reagents were titrated to ensure equivalent staining for each reagent, with final concentrations ranging from 50 to 200 nM. Samples were thawed and washed with CSM. Then cells were barcoded using the transient partial permeabilization approach^41^, in which samples were washed with 0.03% saponin in PBS (PBS-S, Sigma Aldrich) and incubated for 1 hour at room temperature with 200 μL of mass tag barcoding reagents diluted in PBS-S. After three washes with CSM, samples from each plate were pooled together with reference cells in a single reaction tube. Cells were blocked with FcR blocking reagent (Miltenyi Biotec) diluted in Perm-S buffer (Fluidigm) for 10 min at 4°C. After a centrifugation step, cells were resuspended in Perm-S and divided in half to be stained with both antibody panels (600 μL staining volume for up to 10^7 cells) for 1 hour at 4°C. Following three washes in Perm-S and one wash in PBS, cells were fixed with 1.6% formalin (Thermo Fisher) working solution in PBS for 10 min at room temperature in the dark. Pelleted cells were resuspended in 0.4 mL of 0.5 μM iridium-labeled nucleic acid intercalator (Fluidigm) and incubated overnight at 4°C. Prior to CyTOF acquisition, cells were washed sequentially in CSM, PBS, and water, then resuspended at 0.5 × 10^6 cells/mL in Cell Acquisition Solution (Fluidigm) supplemented with 10% EQ Four Element Calibration Beads (Fluidigm). Samples were acquired on a Helios-upgraded CyTOF 2 system, with data collected as individual FCS files. The samples were stained and acquired in 7 batches.

### CyTOF Preprocessing

FCS files from each sample were pre-processed using a semi-automated R pipeline built on the CATALYST framework, which performed file concatenation, bead -based normalization, spillover compensation, sample debarcoding, and batch correction as previously described^42^. Quality control steps included removal of inter-sample doublets based on barcode signal intensities and exclusion of intra-sample doublets based on DNA content (iridium intercalator signal). A spillover compensation matrix was generated and applied to all antibodies in the panel following established protocols^43^.

Cells were gated on cisplatin Pt194 to remove dead cells from the analysis. Following data normalization and batch correction, antibody expression was arcsinh-transformed. Cells were then gated on human CD45 expression to remove non-immune and most murine immune cells. Candidate immune cells were subsequently gated on CD45M-H2Kd to remove any murine immune cells.

### CyTOF Analysis

Cells profiled using the adaptive immune-focused and innate immune-focused panels were analyzed separately. Cells were clustered using a self-organizing map (GigaSOM) with SOM dimensions of 10 x 10, yielding 100 clusters^44^. SOM clusters were subsequently grouped into meta-clusters through consensus clustering (ConsensusClusterPlus, metric = Euclidean distance). Multiple rounds of clustering were performed; initial clustering involved a limited feature set of broad lineage markers (CD3, HLA-DR, CD15, CD20, CD14, CD11c, CD7, CD38, c-KIT, CD45H) and k = 15 metaclusters. Metaclusters were annotated as T cells, NK cells, B cells, granulocytes, other myeloid cells, and non-immune cells, with any remaining non-immune metaclusters being discarded from subsequent analysis. Cells profiled using the adaptive panel and annotated as lymphocytes were subsequently re-clustered using an expanded set of markers and k = 20 metaclusters. Cells profiled using the innate panel and annotated as myeloid cells were also re-clustered using an expanded set of markers and k = 20 metaclusters. Annotation of metaclusters was performed manually by examining the expression of canonical lineage markers.

### IMC Data Acquisition

A condensed panel of 40 antibodies (Table 2), based on the antibody panels selected for CyTOF, was used to analyze formalin-fixed, paraffin -embedded (FFPE) tissue sections from primary tumour and PDX samples. Tissue sections were stained with metal-tagged primary antibodies and metal-conjugated secondary antibodies using a standard protocol, as previously described^45^. Conjugated antibody concentrations were individually titrated on serial FFPE tissue sections of one of the PDX samples chosen at random. Tissue sections were divided into a grid of 1mm squares where at least half of the squares for each sample were randomly selected for imaging. Image acquisition was conducted using the Hyperion Imaging System (Standard BioTools) at ∼1 µm resolution with laser ablation frequency of 400 Hz in a rasterized pattern.

### IMC Analysis

Single-cell masks were generated using the segmentation algorithm DeepCell^46^ and an aggregate of CAIX, PanCK, SMA, CD31, hCD45, mCD45 channels as a membrane channel input image. Protein intensities were quantified on a per-cell basis. Single cell data was corrected for signal spillover between channels using the Bioconductor CATAL YST R package^43^. Cells with a diameter >20 μm or <5 μm were filtered out. The expression of each protein was capped at the 99th percentile of expression, and expression values were then scaled from 0-1 by dividing by this maximum value. Cells were clustered using normalized expression values with the PhenoGraph algorithm and were manually annotated as described in the CyTOF section.

### Statistical Analysis for the Comparison of Distributions

All pairwise combinations of samples were compared using a similarity (Pearson correlation) or distance (Euclidean) metric. Sample pairs were grouped by the metadata of interest into either matching pairs (same metadata label) or non-matching pairs (different metadata labels). Matching and non-matching distributions were compared using Student’s t-test.

## Notes

### Competing Interest Statement

Taconic Biosciences Inc provided the mice used in this study under a sponsored research agreement.
K.A.L. reports receiving honorarium from AbbVie and speaking fees from Virtual Hallway.
H.W.J. has consulted for and received travel and research support from Standard BioTools unrelated to this work.
G.D.B. has equity in Adela Inc. and is an advisor of BioRender.
P.D. is an employee of Taconic Biosciences, Inc.
B.B. has founded and is a shareholder and member of the board of Navignostics, a precision oncology spin-off from the University of Zurich.

### Summary of Updates

Figure 5 was revised to address an error in the labelling of panel 5B

