## Supplementary Figures for "Humanized patient-derived xenografts preserve tumour-specific immune microenvironments"

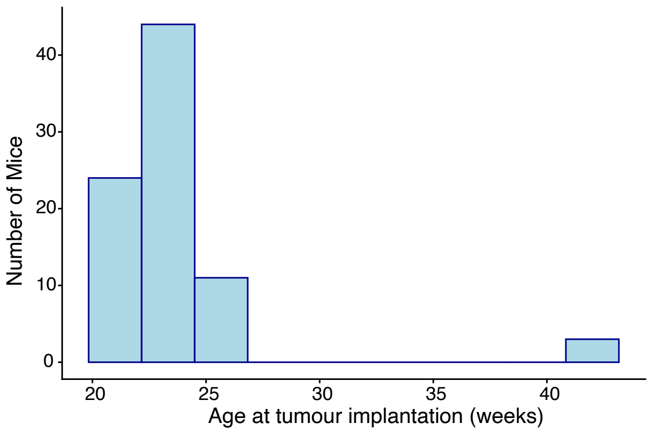


**Figure S1: Age of mice at time of tumour implantation.** Ages are shown in weeks post-humanization which occurs at 4-6 weeks of age. Most mice were implanted with tumour samples between 22 and 26 weeks post-humanization, and four mice were implanted at 42 weeks post-humanization.

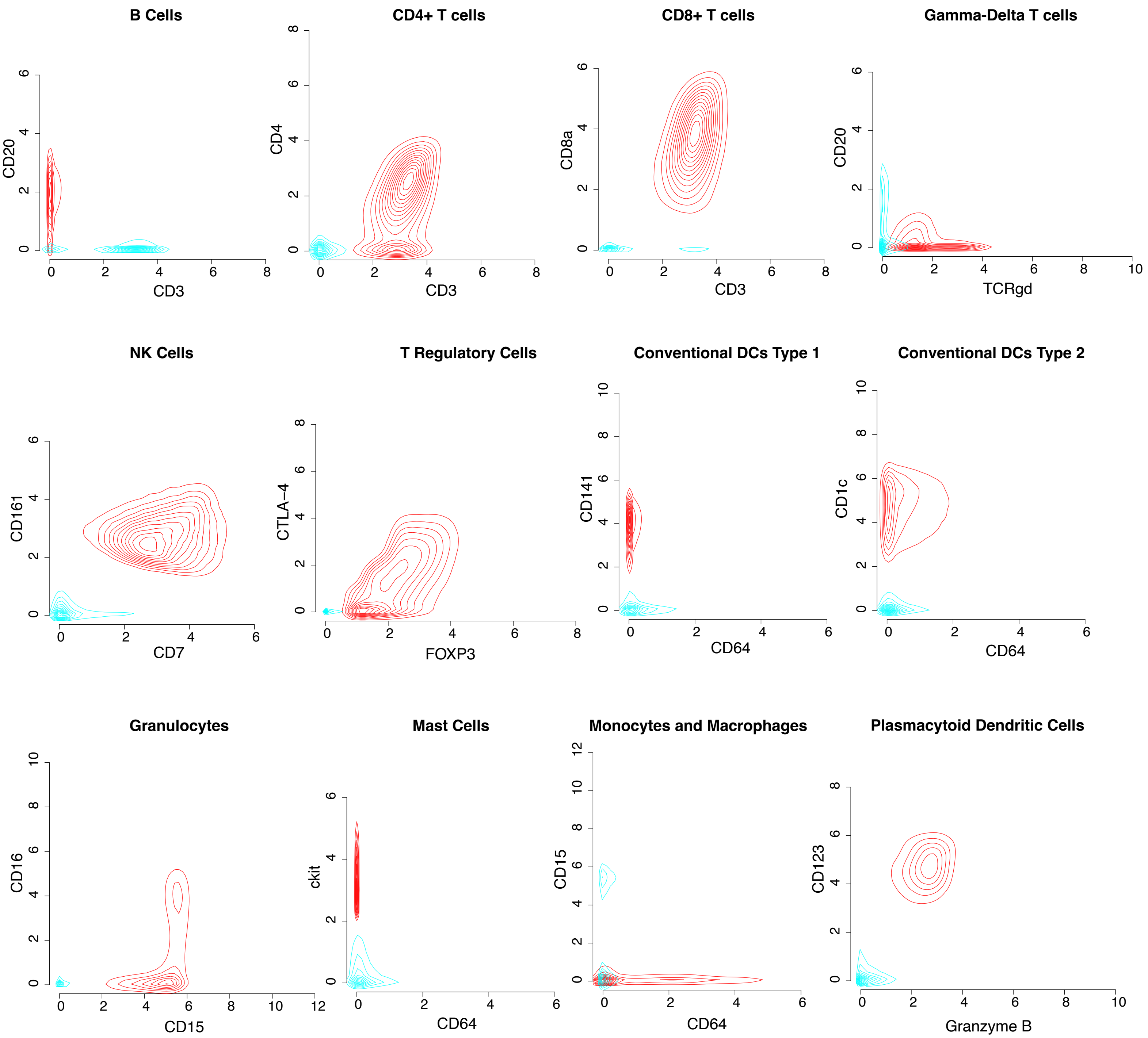


**Figure S2: Annotation strategy for immune cell types.** Density plots demonstrating the normalized expression of two selected markers are shown. In each panel, all cells labelled as the cell type of interest are shown in red, and all other cells are shown in blue. Myeloid cell types of interest (dendritic cells, granulocytes, mast cells, monocytes and macrophages) are compared against all other myeloid cells profiled using the innate panel. Lymphoid cell types of interest (T cells, B cells, NK cells) are compared against all other lymphoid cells profiled using the adaptive panel.

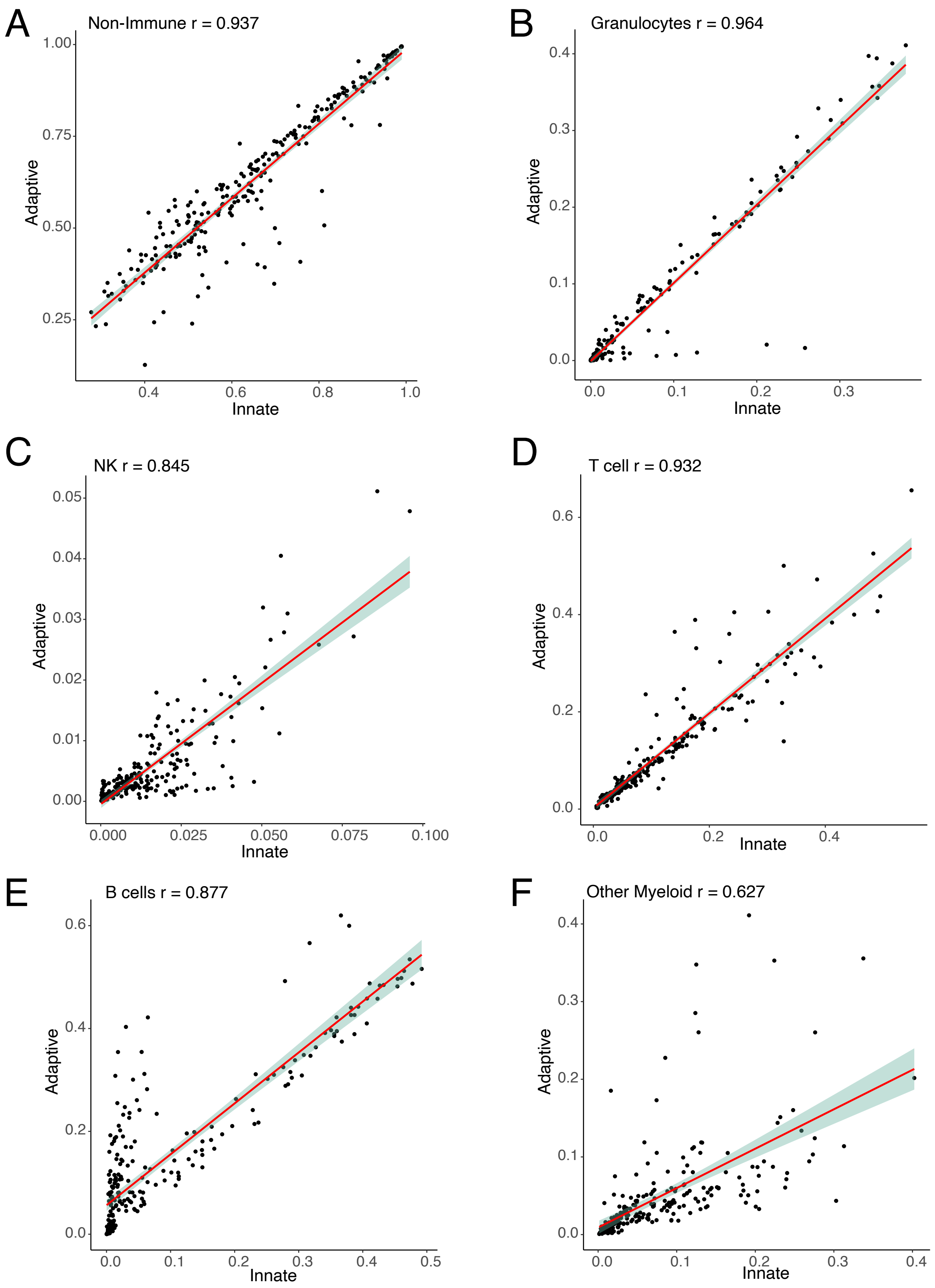


**Figure S3: Cross-panel validation of cell type annotations.** Broad cell type annotations were generate using both the innate and adaptive immune panels for all 249 PDX, bone marrow, spleen, and primary tumour samples using antibodies which overlapped both panels. Cell type proportionsfor each sample annotated in the innate and apative panel datasets are shown as scatter plots, and a linear model is shown in red with standard error for this model shown in teal. Pearson correlation values are shown for each cell type. A) Non-immune cells. B) Granulocytes. C) NK cells. D) T cells. E) B cells. F) Other myeloid cells (monocytes, macrophages, and dendritic cells).

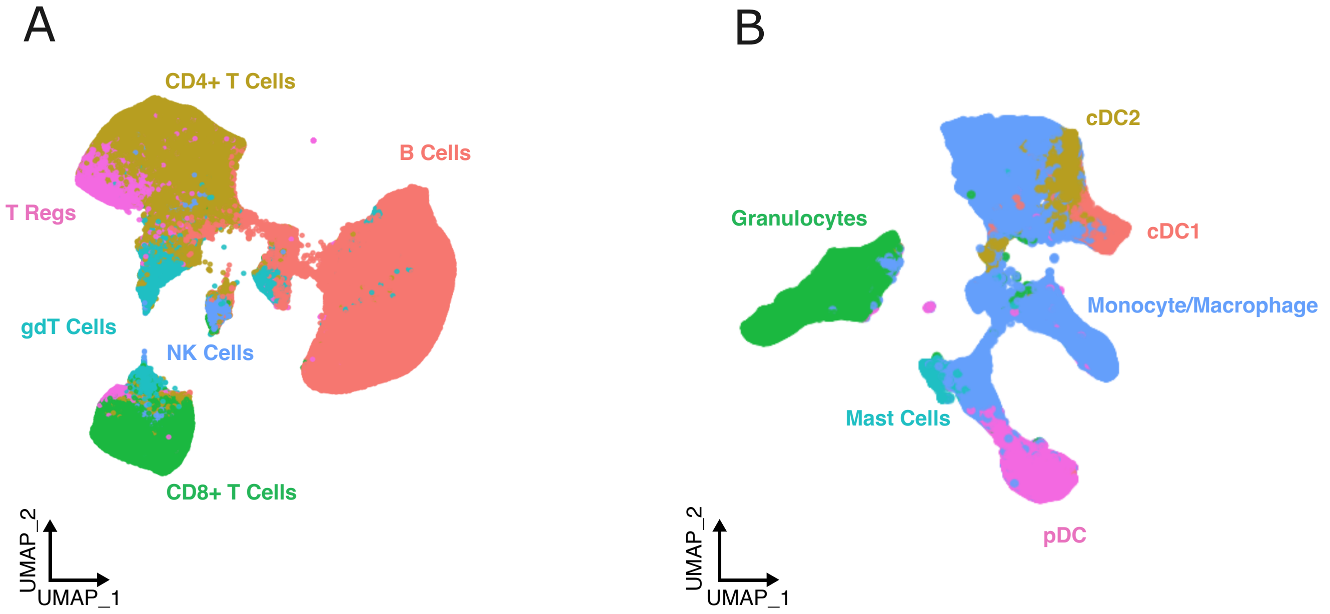


**Figure S4: UMAP representation of annotated immune cell types**. A) UMAP representation of all human immune cells annotated as lymphocytes in the adaptive panel are shown and coloured by cell type annotations. B) UMAP representation of all human immune cells annotated as myeloid cells in the innate panel are shown and coloured by cell type annotations.

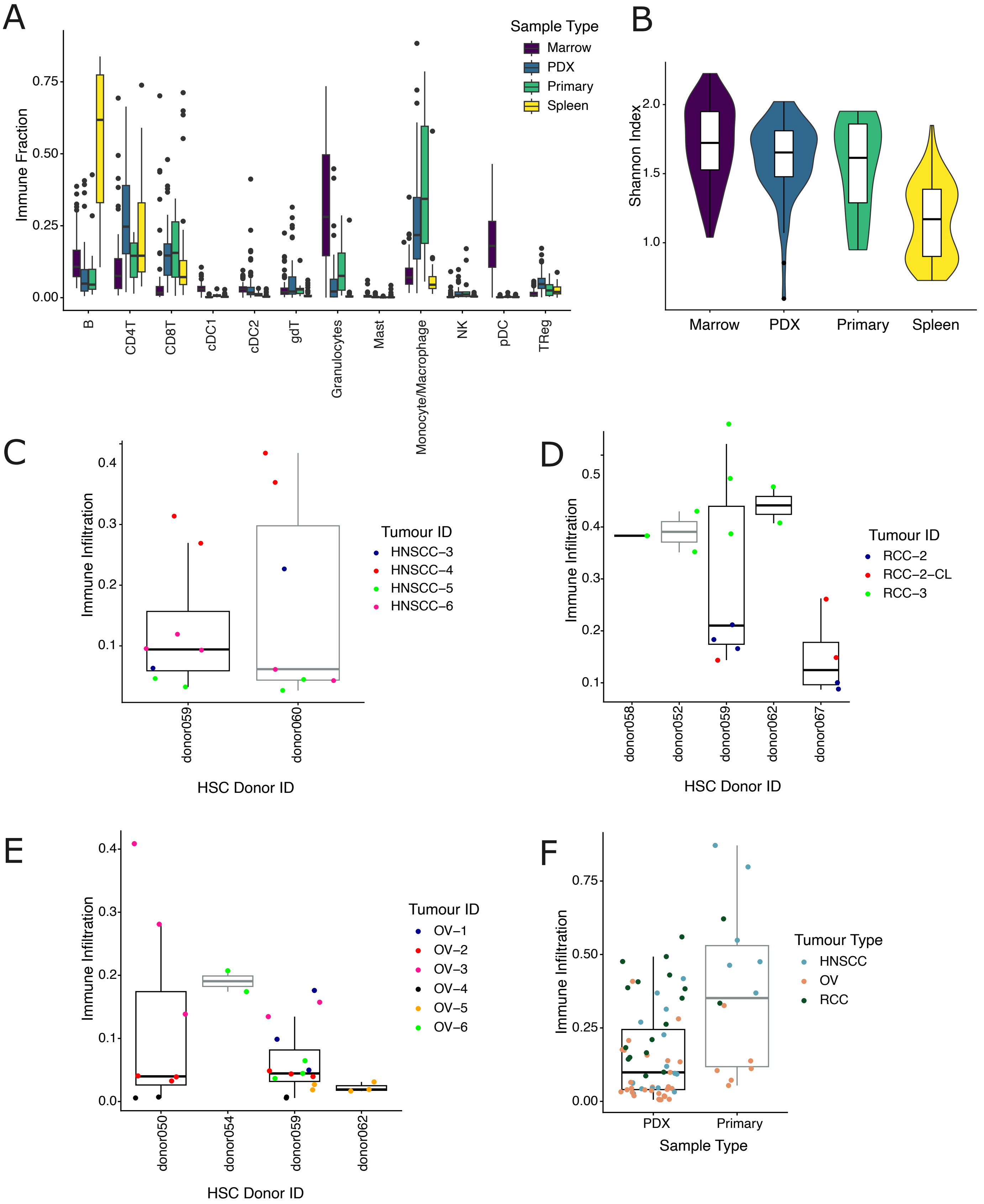


**Figure S5: Immune infiltration and composition across samples and tumour types.** A) Boxplots showing the distribution of human immune cell type abundances across all samples. Samples are grouped and coloured by sample type. Immune cell type abundances are normalized to the total number of human immune cells within each sample. B) The Shannon Diversity Index was calculated using the immune cell type proportions within each sample, and the distribution of these scores across each sample type is shown as a violin plot. C) Immune infiltration within PDX samples was calculated as the number of human immune cells divided by the total number of live cells within each sample. Immune infiltration shown for all PDX originating from HNSCC tumours. Samples are coloured by tumour ID and grouped as boxplots by CD34+ stem cell donor. D) Immune infiltration was calculated as in (C) and shown for all PDX originating from RCC tumours. Samples are coloured by tumour ID and grouped as boxplots by CD34+ stem cell donor. E) Immune infiltration was calculated as in (C) and shown for all PDX originating from OV tumours. Samples are coloured by tumour ID and grouped as boxplots by CD34+ stem cell donor. F) Immune infiltration within primary tumour and PDX samples was calculated as in (C). Samples were only included if data was available for both the primary tumour and at least one matched PDX sample. Boxplots are shown for infiltration values across PDX and primary tumour samples, and individual values are coloured by tumour type.

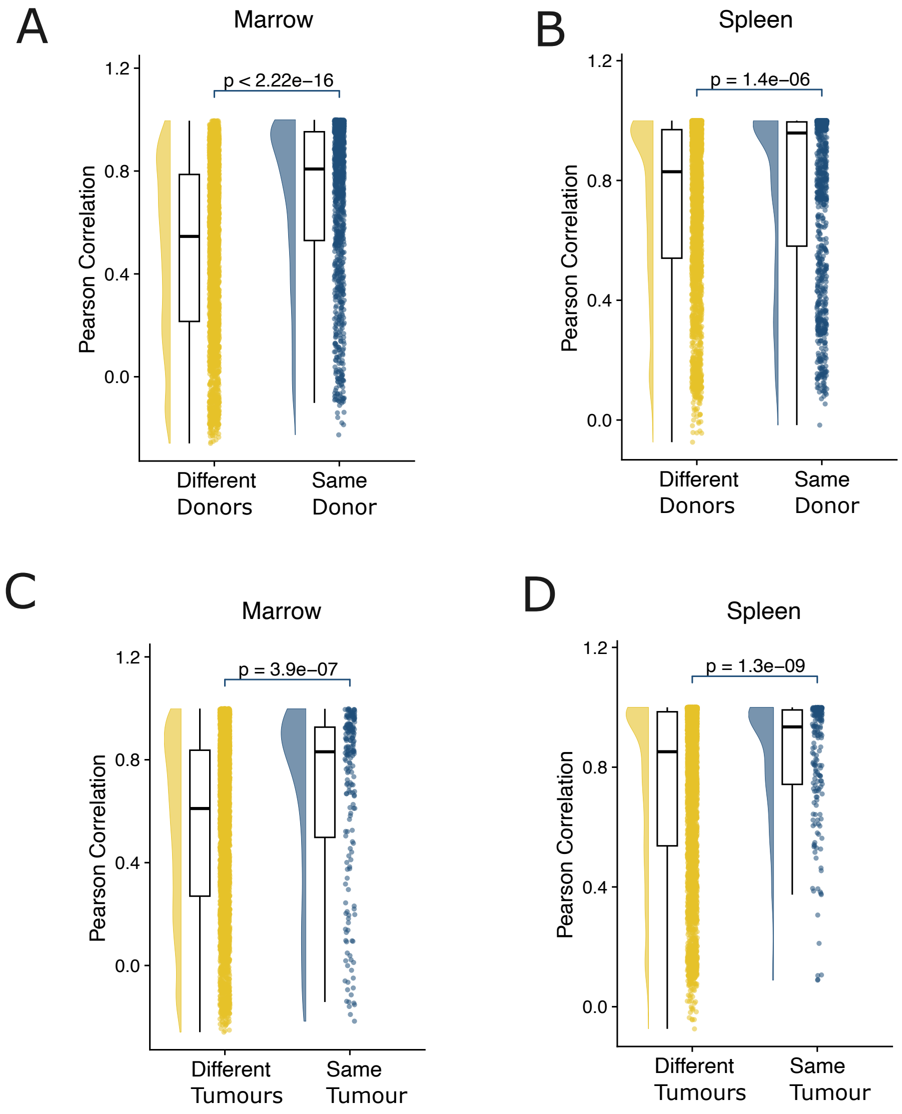


**Figure S6: Systemic immune composition is driven by both HSC donor and engrafted PDX.** A/B) Pairwise similarity of immune cell type composition between all bone marrow (A) or spleen (B) samples. Pairs of samples were grouped depending on if they were humanized using the same stem cells donor (blue), or different donors (yellow). C/D) Pairwise similarity of immune cell type composition between all bone marrow (C) or spleen (D) samples. Pairs of samples were grouped depending on if they were implanted using the same primary tumour (blue), or different primary tumours (yellow). Reported p values indicate the results of Student's t-tests.

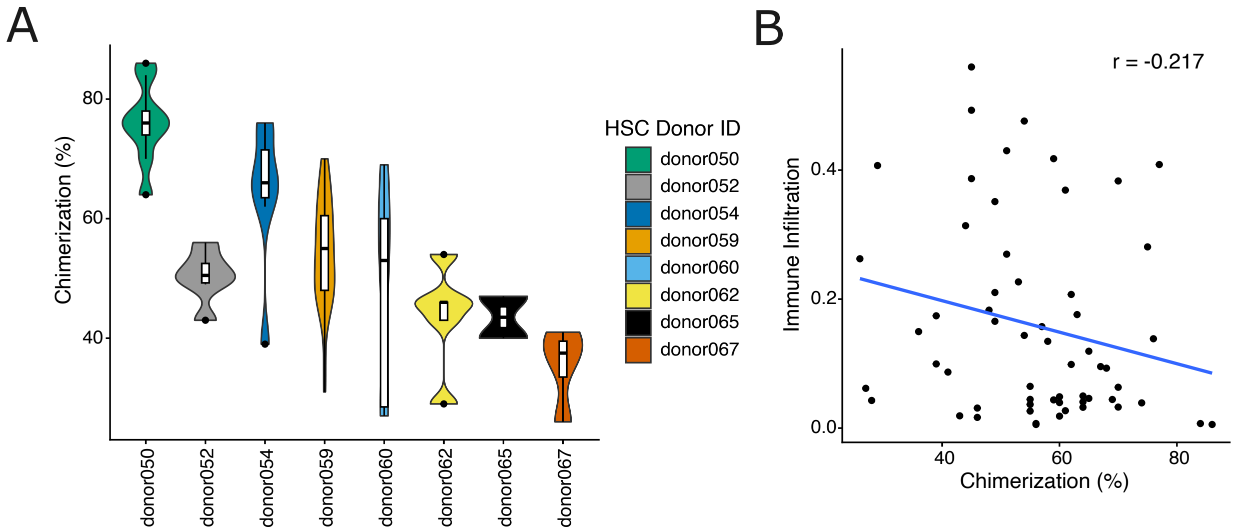


**Figure S7: Immune infiltration in PDX is not driven by donor-dependant chimerization.** A) Blood chimerization values, the proportion of human CD45+ cells in blood samples, were provided by Taconic Biosciences. Chimerization was measured at 10 weeks post-humanization and was determined by flow cytometry by gating on human and murine CD45. Mice were grouped by CD34+ stem cell donor, and violin plots display the distribution of chimerization values within mice humanized using each donor. Data was not shown for donor #058 as only one mouse was humanized using stem cells from this donor. B) Chimerization values for each mouse successfully engrafted with a tumour sample were compared to immune infiltration within each PDX.

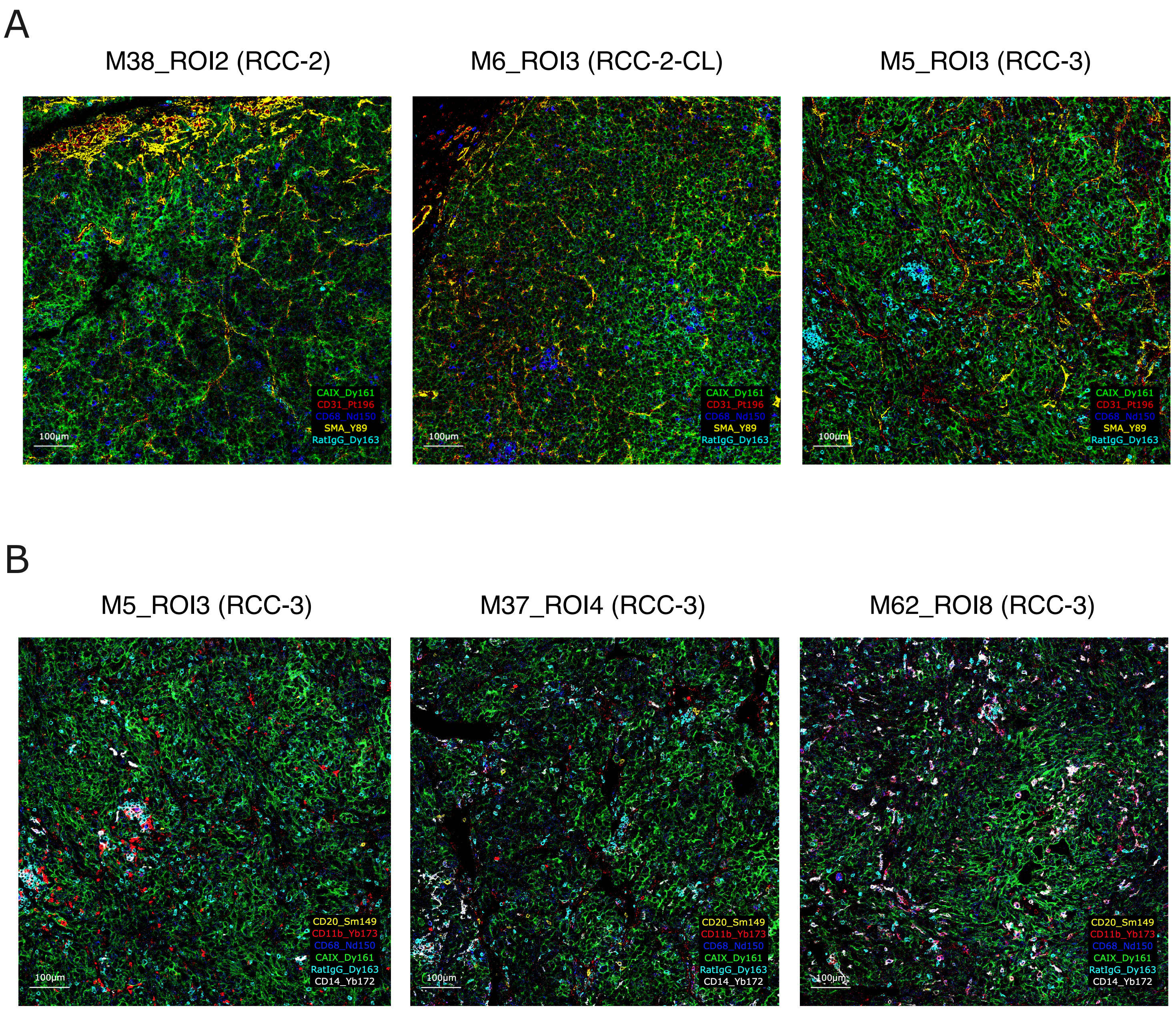


**Figure S8: Validation of immune infiltration in RCC PDX using Imaging Mass Cytometry**. A) One representative Region of Interest (ROI) from PDX originating from RCC-2, RCC-2-CL, and RCC-3 are shown. All samples show high cellularity (green: RCC cells, marked by CA9). Vasculature (red: endothelial cells, marked by CD31), stromal cells (yellow: fibroblasts and pericytes, marked by SMA) myeloid cells (blue: CD68) and T cells (cyan: CD3, measured by secondary antibody for rat anti-CD3) were detected in all samples. B) NK cells are shown in ROIs from three PDX originating from RCC-3. NK cells are marked by CD11b expression and are shown as red (CD11b+, CD68-, CD20-, CD3-, CD14-).

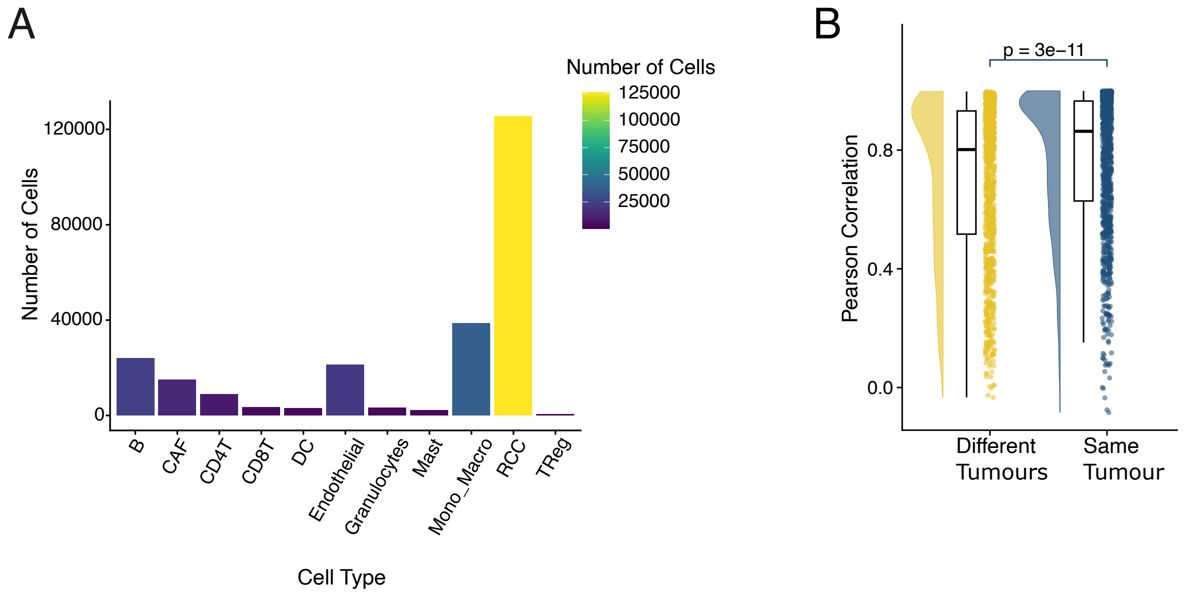


**Figure S9: Cell types annotated in IMC dataset.** A) Barplot showing the total number of cells annotated as each cell type in the IMC samples. B) Pairwise similarity of immune cell type composition was calculated as pearson correlation between all pairs of PDX ROIs. Pairs of ROIs were grouped depending on if they originate from the the same primary tumour (blue), or different primary tumours (yellow). RCC-2 and RCC-2-CL were treated as the same primary tumour for this analysis. Distributions of similarity values were compared using Student's t-test.

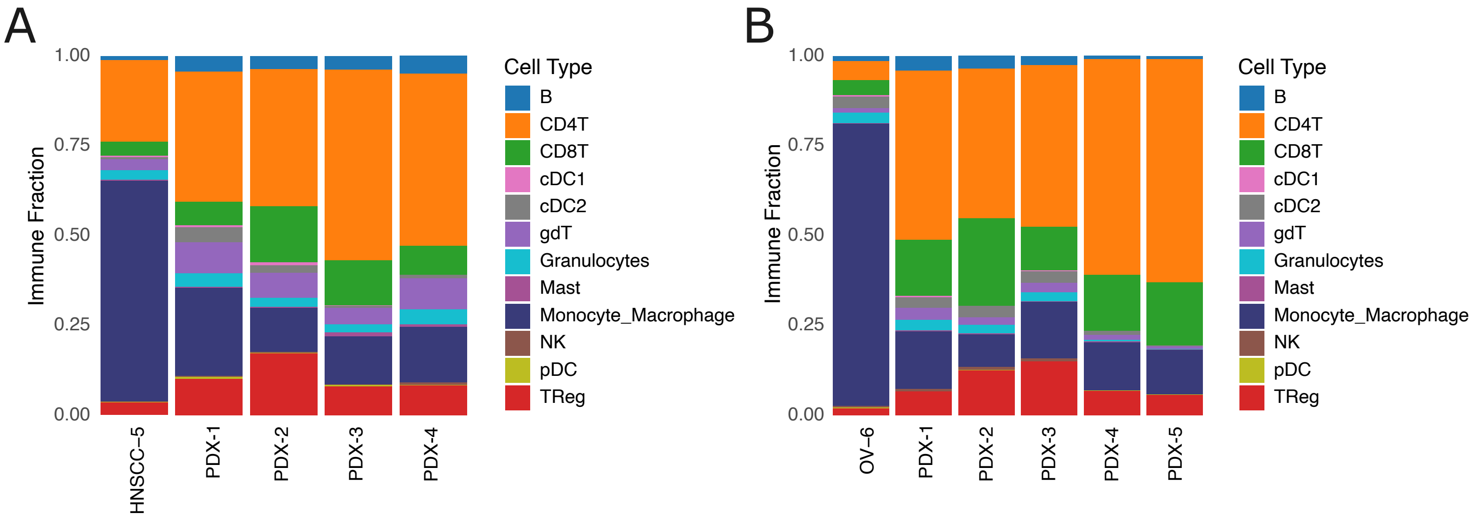


**Figure S10: Case studies of PDX failing to recapitulate myeloid infiltration.** A) Stacked bar plots showing the proportion of each immune cell type in the primary tumour HNSCC-5 alongside all four PDX originating from this tumour. B) Stacked bar plots showing the proportion of each immune cell type in the primary tumour OV-6 alongside all five PDX originating from this tumour.
